# From head to tail: A neuromechanical model of forward locomotion in *C. elegans*

**DOI:** 10.1101/295154

**Authors:** Eduardo J. Izquierdo, Randall D. Beer

**Author notes:** **Author for correspondence:** Eduardo J. Izquierdo. Article submitted to Philosophical Transactions of the Royal Society B. Special Issue: Connectome to behaviour: Modelling *C. elegans* at cellular resolution.

## Abstract

With 302 neurons and a near complete reconstruction of the neural and muscle anatomy at the cellular level, *C. elegans* is an ideal candidate organism to study the neuromechanical basis of behavior. Yet, despite the breadth of knowledge about the neurobiology, anatomy and physics of *C. elegans*, there are still a number of unanswered questions about one of its most basic and fundamental behaviors: forward locomotion. How the rhythmic pattern is generated and propagated along the body is not yet well understood. We report on the development and analysis of a model of forward locomotion that integrates the neuroanatomy, neurophysiology and body mechanics of the worm. Our model is motivated by experimental analysis of the structure of the ventral cord circuitry and the effect of local body curvature on nearby motoneurons. We developed a neuroanatomically-grounded model of the head motoneuron circuit and the ventral nerve cord circuit. We integrated the neural model with an existing biomechanical model of the worm’ s body, with updated musculature and stretch receptors. Unknown parameters were evolved using an evolutionary algorithm to match the speed of the worm on agar. We performed 100 evolutionary runs and consistently found electrophysiological configurations that reproduced realistic control of forward movement. The ensemble of successful solutions reproduced key experimental observations that they were not designed to fit, including the wavelength and frequency of the propagating wave. Analysis of the ensemble revealed that head motoneurons SMD and RMD are sufficient to drive dorsoventral undulations in the head and neck and that short-range posteriorly-directed proprioceptive feedback is sufficient to propagate the wave along the rest of the body.

## Introduction

Behavior is grounded in the interaction between an organism’ s brain, its body, and its environment. How simple neuronal circuits interact with their muscles and mechanical bodies to generate behavior is not yet well understood. With 302 neurons and a near complete reconstruction of the neural and muscle anatomy at the cellular level [1], *C. elegans* is an ideal candidate organism to understand the neuromechanical basis of behavior.

Locomotion is essential to most living organisms. Since nearly the entire behavioral repertoire of *C. elegans* is expressed through movement, understanding the neuromechanical basis of locomotion is especially critical as a foundation upon which analyses of all other behaviors must build. *C. elegans* locomotes in an undulatory fashion, generating thrust by propagating dorsoventral bends along its body. Movement is generated by body wall muscles arranged in staggered pairs along four bundles [2]. The anterior-most muscles are driven by a head motoneuron circuit and the rest of the muscles are driven by motoneurons in the ventral nerve cord (VNC). Although the nematode is not segmented, a statistical analysis of the VNC motoneurons in relation to the position of the muscles they innervate revealed a repeating neural unit [3]. Interestingly, while the repeating neural units in the VNC are interconnected via a set of chemical and electrical synapses, the head motoneuron circuit is largely disconnected from the VNC neural units. Motoneurons in both the head and the VNC circuit have been long postulated to be mechanosensitive to stretch [1, 4, 5], and evidence in support of this has been shown recently for the VNC [6]. Despite all of this anatomical knowledge, how the rhythmic pattern is generated and propagated along the body during forward locomotion on agar is not yet well understood.

A number of computational models of *C. elegans* locomotion have been proposed (see reviews [7, 8, 9]). The model described in this paper differs from previous models in four main ways. First, the current model of the VNC incorporates the analysis of its repeating structure [3]. Second, the current model of stretch-receptor feedback takes into consideration findings regarding the range and directionality of local body curvature on motoneurons [6]. Third, the current model integrates the head motoneuron circuit and the VNC motoneuron circuit within a physical model of the body and environment, such that the forward motion of the model emerges from the undulation of the head, neck, and body. Finally, all current models have assumed specific mechanisms for how the rhythmic movement is generated and propagated, with little systematic exploration of the possibilities.

Here we present a model of forward locomotion grounded in the neurobiology, anatomy, and physics of the worm. The model integrates a head motoneuron circuit based on hypotheses postulated in the original “Mind of the Worm” paper [1] with a model of a repeating neural unit in the VNC based on a statistical analysis of the available connectome data [3]. Motoneurons innervate an anatomically grounded model of the muscles. Stretch receptors are modeled to match recent experimental evidence on the effect of local body curvature on nearby motoneurons [6]. The neuromuscular system is embedded in a model of the physics of the worm’ s body [10]. We used an evolutionary algorithm to explore the space of unknown parameters of the head and VNC motoneuron circuits such that the integrated neuromechanical model matched the speed of the worm during forward locomotion on agar. Analysis of successful solutions suggests that sensory feedback mechanisms in the head motoneurons and the VNC are sufficient to generate and propagate dorsoventral waves to produce forward locomotion behavior. Detailed analysis of the operation of the model sheds further light on the mechanisms that generate and propagate oscillations and leads to a number of experimental predictions.

## Model

### Environment properties

In the laboratory, *C. elegans* is typically grown and studied in petri dishes containing a layer of agar gel. The gel is firm and worms tend to lie on the surface. The locomotion behavior observed under these conditions is referred to as crawling. Worms are sometimes also studied in a liquid medium such as water, leading to a related locomotion behavior called swimming [11]. The experiments in this paper will focus only on agar gel. Given the low Reynolds number physics of *C. elegans* locomotion, inertial forces can be neglected and the resistive forces of the medium can be well-approximated as a linear drag *F* = −*Cv* [10, 12, 13, 14]. Estimated values of the ratio for drag coefficient for nematodes crawling on agar gels vary by as much as an order of magnitude (ranging from 1.5 to 40) in the literature [11, 14, 15, 16, 17]. The tangential and normal drag coefficients for agar used in this model were taken from those reported in [11] and used in the model of the body that this work builds on [10]: *C*_*║*_ = 3.2 ×; 10^−3^ kg·*s*^*−1*^ and *C*_*┴*_ = 128 ×; 10^−3^ kg·*s*^−1^, respectively [10, 11, 12, 14, 18, 19].

### Body model

The model of the body is a reimplementation of the model presented by Boyle, Berri, and Cohen [10]. The worm is modeled in 2D cross-section. This is justified because when placed on an agar surface, the worm locomotes on its side, bending only in the dorsal-ventral plane. The ∼1mm long continuous body of the worm is divided into variable-width discrete segments (Fig. 1A), each of which are bounded by two cross-sectional rigid rods whose endpoints are connected to their neighbors via damped spring lateral elements modeling the stretch resistance of the cuticle and damped spring diagonal elements modeling the compression resistance of internal pressure. The rest lengths, spring constants and damping constants of the lateral and diagonal elements are taken directly from previous work [10], who in turn estimated them from experiments with anesthetized worms [20]. The forces from the lateral and diagonal elements are summed at the endpoints of the rods and then the equations of motion are written for the center of mass of each rod. The full set of expressions for forces are identical to those in [10, 12]. Since each rod has two translational (*x, y*) and one rotational (*ϕ*) degrees of freedom, the body model has a total of 3(*N*_seg_ + 1) degrees of freedom. The current model has *N*_seg_ = 50, so a total of 153 degrees of freedom. All kinematic and dynamic parameters are identical to those used in [10, 12].

**Figure 1.**
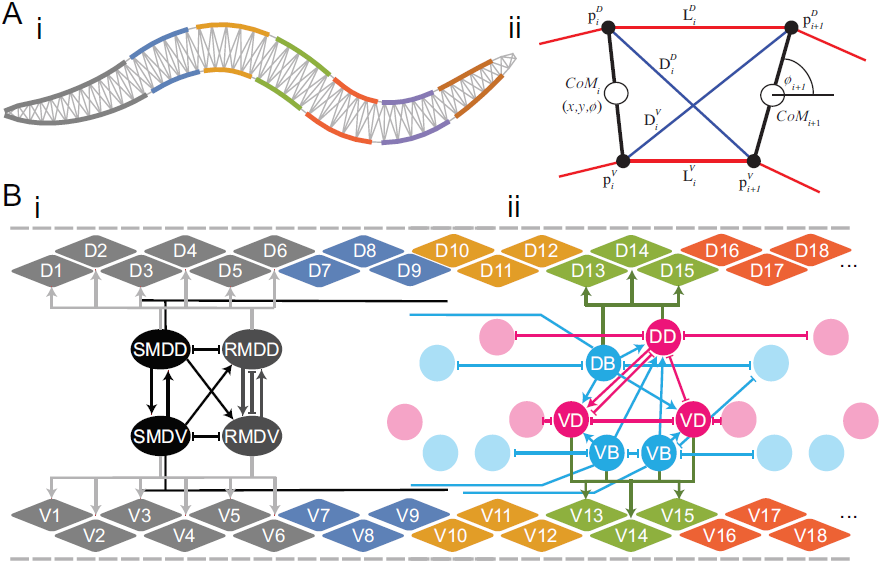
Neuromechanical model. [A] Physical model of the body adapted from [10]: (i) Complete model. Lateral elements are colored according to the muscles they are driven by. Head and neck muscles are driven by the head motoneuron circuit (gray) (see panel B(i)). The rest of the body wall muscles are driven by a series of 6 repeating VNC units (blue, orange, green, red, purple, and brown) (see panel B(ii)). (ii) One of 49 individual segments. Cross-sectional rigid rods (black), damped spring lateral elements (red), damped spring diagonal elements (blue). [B] Neuromuscular model. Dorsal and ventral lateral elements from the physical body represented in gray on the top and bottom, respectively. Dorsal and ventral staggered muscle arrangement. Muscle force is distributed across all lateral elements they intersect. (i) Head neuromuscular unit includes SMD (black) and RMD (gray) motoneurons that connect to muscles on each side. SMD-class neurons receive stretch-receptor input from self and posterior region covered by black process. (ii) One of 6 repeating VNC neuromuscular units, derived from a statistical analysis of the connectome [3]. Each unit includes one dorsal and two ventral B- (blue) and D-class (magenta) motoneurons that connect to muscles on each side. B-class neurons receive stretch-receptor input from anterior region covered by blue process [6]. Circuits include all chemical synapses (arrows), gap junctions (connections with line endings), and neuromuscular junctions.

### Muscles

Body wall muscles in the worm are arranged as staggered pairs in four bundles around the body and are divided into 16 in the head, 16 in the neck and 63 in the rest of the body [2, 21]. These muscles can contract and relax in the dorsoventral plane. Unlike previous work [10], we do not directly associate each discrete lateral element of the body model with a distinct muscle. Instead, muscles are modeled as separate damped springs that lie along the cuticle and their force is distributed across all lateral elements that they intersect (Fig. 1B). This allows us to vary the spatial resolution of the body discretization independently from the number of muscles. It also allows us to accommodate the fact that adjacent body wall muscles overlap one another in *C. elegans*. Since the model is 2D, we combine right and left bundles into a single set of 24 dorsal and 24 ventral muscles, each with twice the strength. Following previous work [10], muscles are modeled as damped springs with activation-dependent rest lengths, spring constants and damping constants, endowing them with simplified Hill-like force-length and force-velocity properties [22]. Muscle activation is modeled as a leaky integrator with a characteristic time scale (*τ*_M_ = 100ms), which crudely agrees with response times of obliquely striated muscle [23]. The muscle activation is represented by the unitless variable 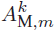 that evolves according to

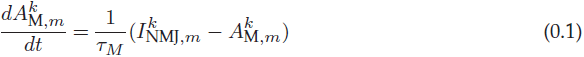

where 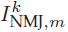 i s the total current driving the muscle. Also following previous modeling work [10] and experimental evidence that electrical coupling between body wall muscle cells only plays a restricted role for *C. elegans* body bend propagation [6, 24], inter-muscle electrical coupling is assumed to be too weak and therefore not included in the model.

### Head motoneuron circuit

In the worm, the head and neck muscles are driven by a set of motoneuron classes that include: bilaterally symmetric RIM, RIV, RMF, RMG, RMH; fourfold symmetric RME, SMB, URA; and sixfold symmetric IL1 [1]. Of these, only four of them (RMD, RME, SMB, SMD) innervate both head muscles and neck muscles; the rest innervate either only the head region (IL1, RMF, RMH, URA) or only the neck region (RIM, RIV, RMG). Given the parallels between SMB and SMD, and between RMD and RME, our model considers only the SMD and RMD motoneurons for the head motoneuron circuit. We used the connectome data to identify the chemical and electrical synapses connecting the two motoneurons and how they innervate head and neck muscles (Fig. 1B(i)). SMD and RMD motoneurons drive head and neck muscles, *m* = [1, 6], according to: 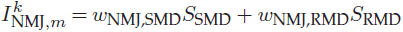. We constrained the sign of their neuromuscular junctions using data from the expression of neurotransmitters: SMD and RMD neuromuscular junctions are both excitatory [25].

### Repeating ventral nerve cord circuit

The rest of the muscles in the body are driven by eight classes of motor neurons: AS, DA, DB and DD innervate the dorsal body wall muscles and VA, VB, VC and VD innervate the ventral muscles. Of the VNC motoneurons, only the B- (DB and VB) and D- (DD and VD) classes have been shown to be involved in forward locomotion, so our model includes them only [6, 26, 27, 28, 29]. As connectome data is incomplete for the posterior half of the worm [1, 30], we relied on a statistical analysis of the motoneurons in relation to the position of the muscles they innervate to model a repeating neural unit along the VNC [3]. When specialized to the B-class and D-class motoneurons, this leads to the circuit architecture shown in Figure 1B (ii). We model 6 such repeating neural units along the VNC, with identical parameters. D- and B-class motoneuron drive body wall muscles posterior to the head and neck, *m* = [7, 24], according to: 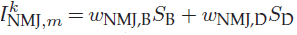. Finally, because the B-class motoneurons are known to be cholinergic and therefore excitatory and the D-class motoneurons are GABAergic and therefore inhibitory [25, 31], we constrain the signs of their neuromuscular junctions accordingly.

### Neural model

Following electrophysiological studies in *C. elegans* [32, 33] and previous modeling efforts [34, 35], all motoneurons were modeled as isopotential nodes with the ability to produce regenerative responses, according to:

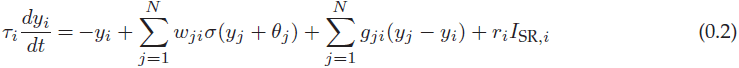

where *yi* represent the membrane potential of the *i*^th^ neuron relative to its resting potential, *τ*_*i*_ is the time constant, *w*_*ji*_ corresponds to the synaptic weight from neuron *j* to neuron *i, g*_*ji*_ corresponds to the conductance between cell *i* and *j* (*g*_*ji*_ *>* 0), and *r*_*i*_ corresponds to the stretch receptor influence to neuron *i*. The model assumes chemical synapses release neurotransmitter tonically and that steady-state postsynaptic voltage is a sigmoidal function of presynaptic voltage [36, 37, 38], *σ*(*x*) = 1*/*(1 + *e*^−*x*^), where *σ*(*x*) is the synaptic potential or output of the neuron (*S*_*i*_). The chemical synapse has two parameters: *θ*_*j*_ is a bias term that shifts the range of sensitivity of the output function, and *w*_*ji*_ represents the strength of the chemical synapse. Electrical or gap junctions between *C. elegans* neurons are common. In line with previous models [35, 38, 39], the model assumes electrical synapses can be modeled as bidirectional ohmic resistances. As we have shown previously [40], this neural model has the capacity to reproduce qualitatively a wide range of electrophysiological properties observed in *C. elegans* neurons [32, 33]. The model can reproduce the passive activity that has been observed in some neurons, like for example, AVA. Through the increase of the strength of the self-connection (*>*4, see [41]), the model is also capable of reproducing the bistable potentials found in some neurons, like, for example RMD [33].

### Stretch receptors

Mechanosensitive stretch receptor channels have long been postulated to exist in motoneurons. There is evidence that supports their existence in interneurons [42, 43], as well as more recently in VNC motoneurons as well [6].

In the head motoneuron circuit, the SMD class has long undifferentiated processes that are distal to the regions where neuromuscular junctions are situated, before they eventually terminate, which have been postulated to be stretch sensitive [1]. We model SMD-class motoneuron stretch receptors as a relatively long-range connection spanning the neck muscles and the muscles associated with the first VNC neural unit (*m* = [4, 9]) (Fig. 1B(i)), with the effect that the head and neck regions bend in the same direction and shortly after the bending of the neck and anterior-most body region. The stretch-receptor current for the SMD-class motoneuron sums over contributions from a total of 14 mechanical elements associated with those muscles,

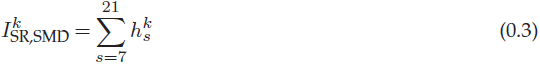

In the repeating neural units of the VNC, the B-class is one motoneuron that has been postulated to mediate stretch-receptor feedback from the body. The long undifferentiated processes running posteriorly have led previous models to assume stretch receptors covered a wide range of muscle cells and that proprioceptive information traveled anteriorly. However, more recent experimental work demonstrated that the effect has a much shorter range than previously assumed and is in fact directed posteriorly, since the activity of each VB and DB motoneuron is activated by ventral and dorsal bending of a more anterior region, respectively [6]. In light of this evidence, we model B-class motoneuron stretch receptors as short-range connections from the lengths of anterior muscles to the immediately posterior B-class motoneurons, with the effect that posterior body regions are encouraged to bend in the same direction and shortly after the bending of a neighboring anterior region (Fig. 1B(ii)). The stretch-receptor current for the B-class motoneuron in unit *n* on the *k*th side,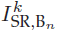, sums over contributions from the *S* = 6 mechanical elements anterior to the anterior-most muscle that neuron innervates (*S*_0,*n*_):

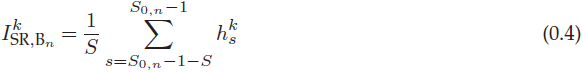

The proposed mechanosensitive channels in these processes respond to the changes in length associated with body bending. In line with previous work [10], stretch receptors are modeled as a weighted linear function of muscle length,

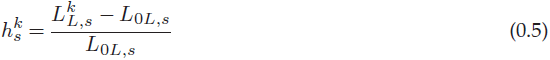

where *L*_0*L,s*_ is the segment rest length and 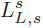 is the current length of the *k*th side (dorsal/ventral) of the *s*th segment. In line with recent findings [6], we allow the stretch receptor conductance to generate a depolarizing response to compression and a polarizing response to stretch, relative to the local segment resting length.

### Numerical methods

The model was implemented in C++ and was solved by Euler integration with a 1ms step. The code for the model and the evolutionary algorithm can be found at https://github.com/edizquie/RoyalSociety2018.

### Evolutionary algorithm

Unknown model parameters were adjusted using a real-valued evolutionary algorithm. A search begins with a random population of genetic strings that encode the unknown parameters of the neural circuit model. Each individual is then assigned a fitness based on the quality of its locomotion performance. Individuals are then selected to serve as parents for the next generation with a probability related to their fitness. From the selected parents, a new generation of children are then produced by randomly swapping portions of two parents (crossover) and making a small modification to the values of the resulting array with values drawn from a Gaussian distribution (mutation). Once a new population has been constructed in this manner, the entire process of evaluation, selection and reproduction repeats until the population converges on highly fit individuals.

A naive parameterization of our model would contain over 400 muscle, neural and stretch receptor parameters. However, it makes little sense to work directly with such a large set of unconstrained parameters. Instead, we imposed a variety of symmetries on the model in order to reduce the number of parameters. We assumed: (a) dorsal/ventral symmetry in the parameters where possible; (b) that the parameters in each VNC neural unit were identical; and (c) that neurons from the same class had identical parameters. Altogether, the model has 30 free parameters. 4 Biases, 4 time-constants, 4 self-connections, and 4 neuromuscular junctions, one for each motoneuron class (class). 2 stretch-receptor gains for SMD and B stretch-receptors. In the head motoneuron circuit, weights for: 3 chemical synapses (synapses between SMD motoneurons, synapses from SMD to RMD motoneurons, synapses between RMD motoneurons); 2 gap junctions (synapse between RMD motoneurons, synapses between SMD and RMD). In the repeating VNC neural unit, weights for: 3 chemical synapses (synapses from B- to D- motoneurons in the same side, synapses from B- to D- motoneurons on opposite sides, and synapse between D- motoneurons); 1 gap junction within the unit (synapse between D- class motoneurons); 3 gap junctions across units (synapses across neighboring D-class motoneurons, synapses across neighboring B-class neurons, synapse on neighboring B-class neurons on opposite sides). Some parameters were constrained to match experimental observations. Specifically, the self-connection for RMD was constrained to> 4 to force the model neuron to be bistable as observed experimentally [33] and neuromuscular junctions were constrained to be positive or negative depending on data from the expression of their neurotransmitters.

In order to evaluate the fitness of a solution, we measured the locomotion efficiency of the entire neuromechanical model. Specifically, we optimized model worms to match the worm’ s average velocity on agar, by maximizing

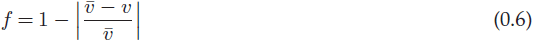

where *v* is the average velocity of the model worm measured over 50 simulated seconds and 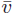 is the average velocity of the worm (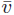 = 0.22mm/sec, based on the ranges reported experimentally [44, 45, 46, 47]). We measure the average velocity of the model worm by calculating the Euclidean distance from the location of the center of the model worm’ s body at the beginning of a trial to the location of its center at the end of the trial.

## Results

### Evolving locomotion

#### Model reliably evolves to match the worm’ s speed

In order to identify circuits that produced forward locomotion, we ran the evolutionary algorithm 100 times using different random seeds. The fitness of the model worm was evaluated to match the worm’ s average velocity on agar (*v* = 0.22mm/sec), based on the ranges reported experimentally [44, 45, 46, 47]. From each evolutionary run, we selected the best individual. As our main interest was to identify networks capable of closely matching the worm’ s behavior, we focused only on the highest performing subset of solutions, namely those networks having a fitness score of at least 0.95 (*n* = 46). All solutions in this subset generated forward thrust by means of a dorsoventral undulation of the body. All further analysis was limited to this ensemble of solutions.

#### Solutions in the ensemble reproduce characteristic features of worm’s movement

The behavior of the models match not only the speed of the worm, but also the overall qualitative kinematics of forward movement. When placed on agar, the models in the ensemble initiate dorsoventral oscillations in the head and propagate them posteriorly, generating thrust against their environment, propelling themselves forward (see movie in Supplementary material). The models can do this robustly, regardless of the initial state and posture of the worm, including from a straight posture. The movement of the model worms resembles the worm’ s characteristic frequency and its wavelength on agar. The ensemble of high-performance solutions locomote with frequencies in the range [0.34, 0.43] and wavelengths in [0.70, 0.96], which are within the range of what has been described in the literature: [0.25, 0.58] [11, 44, 46, 47, 48, 49] and [0.45, 0.83] [11, 44, 46, 47, 48, 49, 50, 51, 52, 53], respectively. That the solutions in the ensemble reproduce characteristic features of the worm’ s movement that they were not evolved to match suggests the model captures fundamental principles of the neuromechanical basis for the behavior in the worm.

### Individual Solution

In order to understand how oscillations are generated and propagated in the model worms, we first consider the operation of one representative individual solution in detail (model parameters in Supplementary material).

#### Head motoneuron circuit can generate oscillations using stretch-receptor feedback

Unlike previous models, the current model makes no explicit a priori assumption about where oscillations should originate. As with the worm, curvature along the body of the model worm over time during forward locomotion suggests the oscillation originates in the head and is propagated posteriorly (Fig. 2A). In order to test whether the head motoneuron circuit can generate oscillations, we silenced motoneurons in the VNC. Even in the absence of oscillatory activity in the VNC, the head could still oscillate (Fig. 2B).

**Figure 2.**
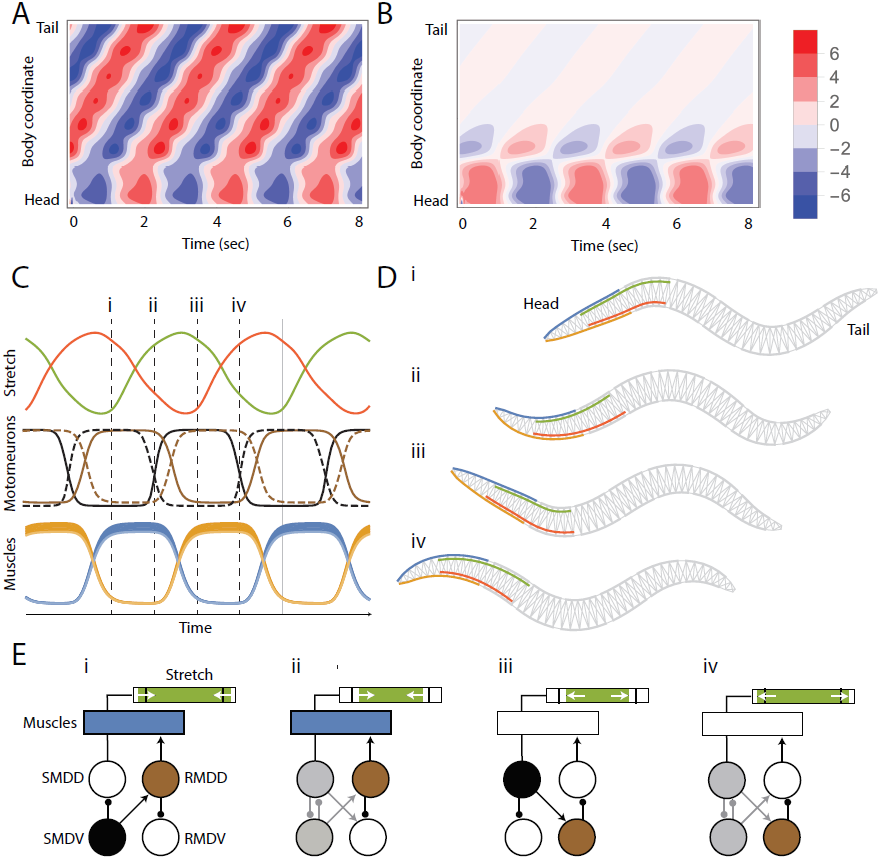
Oscillations in the head motoneuron circuit. [A] Kymogram during normal operation: Oscillation originates in the head and travels posteriorly. [B] Kymogram with VNC motoneurons silenced: Dorsoventral bends persist in head and neck. [C] Traces from stretch receptors, motoneurons, and muscles. Green/red traces dorsal/ventral stretch receptors. Black/brown traces SMD/RMD neural activity. Solid/dashed lines represent dorsal/ventral motoneurons. Blue/orange represents muscle activity from the 6 head and neck dorsal/ventral muscles. Activity is cyclic so four points are chosen in the cycle: i-iv. [D] Postures at the four instances of time selected in panel C. Dorsal/ventral head and neck muscles represented in blue/orange. Dorsal/ventral undifferentiated processes providing stretch information represented in green/red. [E] Mechanics of oscillation. Green bar represents amount of stretch/contraction in the dorsal undifferentiated process with respect to resting state (black vertical line). White arrows represent whether the process is stretching or compressing. Blue rectangle represents the dorsal head and neck muscles. Only dorsal muscles and stretch receptors are shown. The circles below represent the motoneurons. Muscles/neurons are filled in with color when they are contracted/activated and no color when they are relaxed/inactivated. The shade of gray represents the SMD neuron mid-activation. SMD motoneurons are shown in black and RMD motoneurons are shown in brown. Synapses appear only when they are in use.

During regular forward locomotion, motoneurons in the head circuit of the model worm oscillate (Fig. 2C). How are these oscillations generated? To address this question, we first silenced stretch-receptors feedback in the head. When we silence stretch-receptor feedback to the head motoneuron circuit, the neural oscillations in the head motoneuron circuit cease. Therefore, despite the capacity of the head motoneuron circuit to generate intrinsic network oscillations, the model worm produces oscillations robustly through stretch-receptor feedback. Such a reflexive pattern generator hypothesis for oscillations in the head motoneuron circuit had only been considered in two other models previously [49, 54]. We examine the differences between previous models and the current model in detail in the Discussion.

In order to understand how the oscillation is generated through stretch-receptor feedback, we consider the neural traces of the head motoneurons, stretch-receptor feedback, muscle activation, and posture of the body over time during a full cycle of locomotion (Fig. 2C-E). At the start of a cycle (stage i), the head and neck sections are straight (Fig. 2Di), SMD’ s undifferentiated process is stretched and compressing, SMDD is off and RMDD is on (Fig. 2Ei). RMDD activates the dorsal head and neck muscles and inhibits the contralateral RMDV motoneuron. As a result, the dorsal head and neck segments contract, while the ventral segments expand, leading to a dorsal head sweep, and the start of stage ii (Fig. 2Eii). Dorsal contraction in the anterior region of the body leads to activation of the SMDD motoneuron through stretch-receptor feedback, which inhibits SMDV and excites RMDV, causing RMDV to deactivate. Deactivation of RMDV allows the dorsal muscle to begin to relax, and leads to stage iii (Fig. 2Eiii). Stage iii is dorsoventrally symmetric to stage i: the posture of the head and neck are straight, but the state of the neurons are flipped in the dorsalventral dimension. SMDD is now on, and as a result SMDV is off and RMDV is on, which results in RMDD being off. This means the ventral muscles are contracting and the dorsal muscles are relaxing, leading to a ventral head sweep, and the start of stage iv (Fig. 2Eiii). Stage iv is dorsoventrally symmetric to stage ii: the relaxing dorsal segments leads to inactivation of SMDD, which ceases to inhibit SMDV and ceases to excite RMDV. Again together re-activation of SMDV and re-inactivation of RMDV lead to the re-activation of RMDD, which leads to the dorsal muscles contracting again, and the head and neck posture to get back to straight.

#### Oscillatory wave can be propagated posteriorly through stretch receptor feedback and without bistable motoneurons

How is the oscillation that is generated in the head then propagated posteriorly to produce the sinusoidal traveling wave responsible for forward thrust in the model worm? In order to understand the operation of the repeating VNC circuit, we start by simplifying the circuit architecture. Although neural traces suggest B- and D- class motoneurons are active, silencing D-class motoneurons does not affect locomotion performance. Silencing B-class motoneurons or removing the stretch-receptor feedback causes the propagation of the wave to cease. This suggests we can simplify this circuit to only the B-class motoneurons for analysis of the wave propagation. With this simplification, the operation of the VNC circuit is straightforward. As the length of the segment anterior to the neural unit compresses, the stretch receptor excites the motoneuron, activating the muscle, and ultimately causing the contraction of its own segment. We can see this on the ventral side in stages ii and iii, and on the dorsal side on stages iv and i (Fig. 3, panels B and C). Therefore, B-class motoneurons with input from stretch-receptors with information about the length of the anterior regions of the body are the primary drivers of the propagation of the rhythmic wave in this solution. Interestingly, B-class motoneurons are not bistable. Therefore, provided the directionality of stretch-receptor feedback shown in [6], bistable motoneurons are not essential for sustaining proprioceptively driven dorsoventral undulations in the model. However, there are two other components that play roles in the propagation of the wave: the inter-unit gap junctions, and the mechanics of the body. We characterize the contribution of each component individually next.

**Figure 3.**
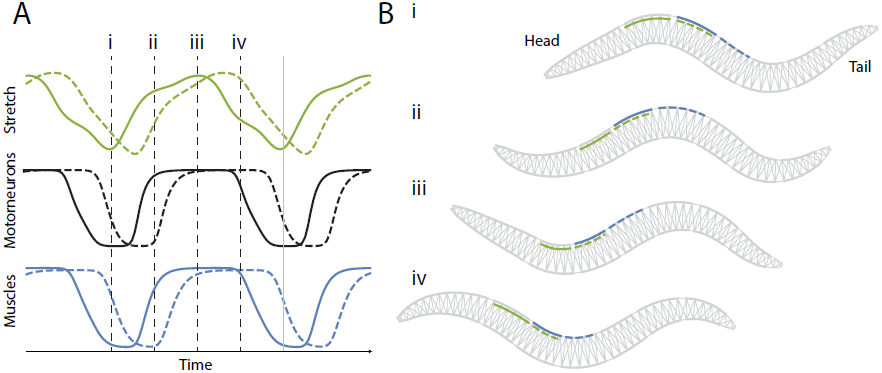
Wave propagation through stretch reception. [A] Traces from the dorsal stretch receptors (green), DB motoneurons (black), and dorsal muscles (blue) in two neighboring VNC neural units: second (solid) and third (dashed). The activity is cyclic so the same four unique points used for Figure 2 were chosen to analyze the wave propagation: i-iv (vertical dashed lines). [B] Worm postures at the four instances of time selected in panel A. The second VNC neural unit receives dorsal stretch receptor input from the solid green region and innervates the muscles in the solid blue region. The third VNC neural unit (posterior to the second), receives dorsal stretch receptor input from the dashed green region and innervates the muscles in the dashed blue region.

#### Inter-unit gap junctions dampens curvature

In the model worm, the propagation of the oscillatory wave from the head to the first unit of the VNC occurs through stretch receptors exclusively, as there are no direct synapses between the head motoneuron circuit and the VNC motoneurons. However, the rest of the VNC units are interconnected by electrical gap junctions between neighboring B-cells (see Fig. 1B). What role do the gap junctions play in transferring the wave posteriorly from the first VNC to the rest of them? When we silenced gap junctions between neighboring units, the wave still travelled posteriorly. Interestingly, the amplitude of the dorsoventral curvature increased by 22%. This suggests gap junctions are responsible for dampening the strength of the curvature. This dampening is functional for forward locomotion: without inter-unit gap junctions, the speed of the model worm dropped to 88.7% of its original speed. In terms of the worm’ s movement, although the frequency of the oscillations remained relatively unaffected, the wavelength became smaller: from 0.81 to 0.68. Altogether, this suggests that when the wave travels through stretch-receptor feedback alone, it travels fast, and the gap junctions between neighboring units act to dampen the wave through tighter communication with the motoneurons. Altogether, while the inter-unit gap junctions play a role in the propagation of the wave, they are not essential for producing forward movement.

#### Wave also propagates through the mechanical body

One of the benefits of a neuromechanical model is that we can study the effect of the mechanical properties of the body on the operation of the behavior. So what role does the body mechanics play in the wave propagation in the model worm? In order to address this question, we silenced the motoneuron activity of each neural unit individually, including the incoming stretch receptor feedback, and the gap junction connections with the unit anterior and posterior to them. Despite the silencing of entire neural units in the VNC, the model worm could still move forward (Fig. 4A). That is, the model worm can recover the traveling wave in the absence of the ventral nerve units from the passive propagation of the wave through the mechanical body. This is because mechanical curvature in one area of the worm forces curvature of neighboring segments. The combination of stretch-receptor feedback and passive mechanical propagation is sufficiently strong that even entirely disabling two adjacent VNC neural units does not impair the ability of a posterior VNC unit from picking up the remains of the traveling wave and re-establishing regular dorsoventral undulations (Fig. 4B).

**Figure 4.**
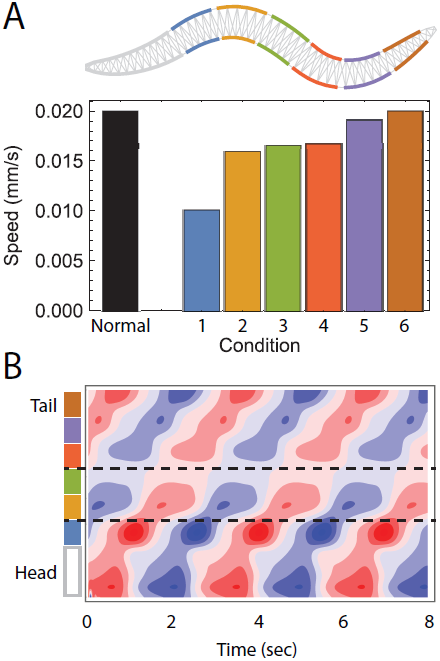
Role of biomechanics in the propagation of the wave and locomotion. [A] Speed of the worm as a result of silencing entire VNC neural units. Color coding according to the region of the body those neural units affect. Black represents the speed of the model worm under normal conditions. Propagation of the wave does not depend entirely on stretch-receptor feedback and neural activity in general. [B] Example kymogram of movement while two VNC neural units (2 and 3) have been silenced. Despite the lack of neural activity, and the lack of network oscillators in the tail, there are oscillations in the head and tail.

### Ensemble of solutions

In the individual solution analyzed in detail, the model moved forward in the absence of an intrinsic network oscillator in either the head motoneuron circuit or the VNC. Instead, oscillations were generated and propagated using stretch-receptor feedback with mechanical propagation playing a substantial role and electrical coupling playing a secondary role. In this section, we analyze how representative that solution is with respect to the rest of the solutions in the ensemble.

#### Wave originates in the head via stretch-receptor feedback or intrinsic network oscillators

All solutions in the ensemble come to a stop when head motoneurons are silenced (orange, Fig. 5A). Yet, when VNC motoneurons are silenced, the head continues to oscillate (green, Fig. 5B), moving forward at a fraction of the speed (green, Fig. 5A). Therefore, in all solutions, the head motoneuron circuit generates oscillations that are used for moving forward. In 40 of the 46 solutions in the ensemble, oscillations in the head ceased when we silenced stretch-receptor feedback to the head motoneuron circuit (red, Fig. 5B). The remaining 6 solutions generate intrinsic network oscillations in the absence of stretch-receptor feedback. These oscillations were sufficient to drive regular forward locomotion (red, Fig. 5A). This suggests the architecture of the head motoneuron circuit can generate oscillations to drive forward locomotion equally well either through intrinsic network oscillations or through stretch-receptor feedback. In both types of solutions, both SMD and RMD motoneurons were essential for producing forward movement throughout the ensemble.

**Figure 5.**
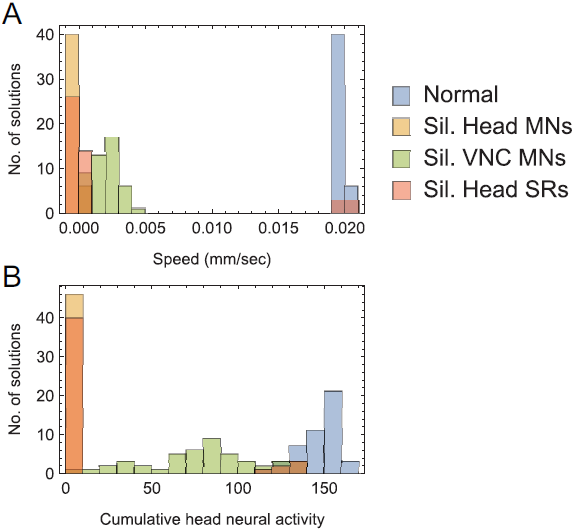
Operation of the head motoneuron circuit in the ensemble of solutions. Distribution of speed [A] and magnitude of change in neural activity in head motoneurons [B] of all model worms in the ensemble under different conditions: Normal locomotion (blue), when head motoneurons are silenced (orange), when VNC motoneurons are silenced (green), when head stretch-receptor feedback is silenced (red).

#### Oscillatory wave is propagated posteriorly through stretch receptor feedback

The way the wave is propagated posteriorly in the ensemble of solutions resembles closely that of the model worm analyzed individually. In order to analyze wave propagation in the ensemble of solutions, we silenced the main components of the VNC while measuring the speed of the worm as well as the average magnitude of the dorsoventral bends along the VNC region of the body (Fig. 6). We summarize the main results ahead. First, the B-class motoneuron is essential for forward locomotion in all solutions. Silencing B-class motoneurons eliminates dorsoventral rhythmic patterns along the body and results in model worms coming to a full stop. Second, B-class motoneurons did not evolve to be bistable in any of the solutions. Therefore, bistable motoneurons are not essential for sustaining proprioceptively driven dorsoventral undulations in the model. Third, silencing stretch-receptor feedback input into the B-class motoneurons also eliminates dorsoventral rhythmic patterns along the body and results in model worms coming to a full stop. Therefore, as with the model worm analyzed individually, stretch receptor feedback is essential for propagating the wave posteriorly. Fourth, in 41 of the 46 solutions in the ensemble, the D-class motoneuron was not essential for forward locomotion. In these solutions, silencing the D-class motoneurons does not affect speed or dorsoventral bends. In the remaining 5 solutions, the D-class is involved in contralateral inhibition and is essential for wave propagation. Fifth, the inter-unit neighboring gap junctions play a minor role in the propagation of the wave. Removing neighboring gap junction augments the strength of the curvature, yet this increase in curvature leads to impaired movement. Finally, the biomechanics of the body alone plays a substantial role in propagating the wave posteriorly. Silencing entire neural units in the VNC does not entirely disrupt propagation of the wave posteriorly. Although silencing entire neural units affects the speed, the model worms still move forward. As with the solution analyzed individually, impairing anterior units has a larger effect than impairing posterior units.

**Figure 6.**
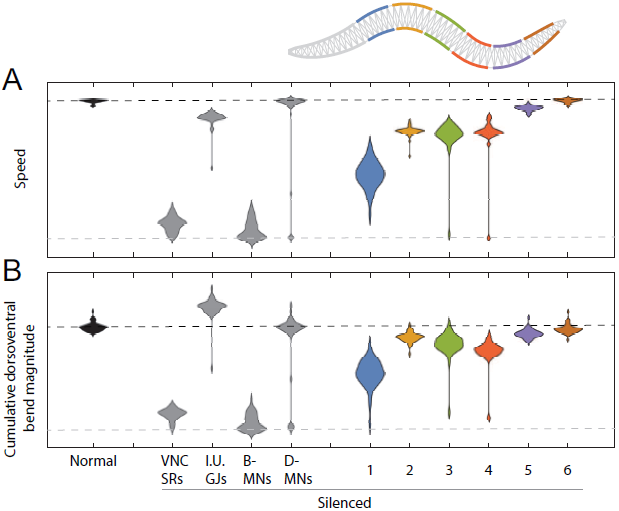
Operation of the VNC in the ensemble of solutions. Distribution of speed [A] and magnitude of dorsoventral bends [B] of all model worms in the ensemble under different conditions: Normal locomotion (black), when VNC stretch-receptor feedback, interunit gap junctions, B-class, and D-class motoneurons are silenced independently (gray), and when an entire neural unit is silenced (colored according to position along the body). The black dashed lines represents the value expected of a normally moving model worm; the gray dashed line represents the value expected of a non-moving model worm.

## Discussion

We have presented a fully integrated, biologically and physically grounded model that accounts for *C. elegans* locomotion on agar that takes into consideration the head motoneuron circuit and the ventral nerve cord motorneuron circuit. The model was motivated by findings regarding the range and directionality of local body curvature on motoneurons [6] and the statistical analysis of the repeating structure of the VNC [3]. With these biological constraints provided, we used an evolutionary algorithm to systematically explore the space of possibilities for generating locomotion. We discuss ahead key insights revealed from the analysis of evolved solutions and related work.

We have demonstrated that a model of the head motornerneuron circuit with SMD and RMD alone is sufficient to generate oscillations that can drive dorsoventral undulations in the head and neck. Analysis of the variations in the ensemble of solutions revealed two possible mechanisms: an intrinsic network oscillator and an oscillator driven by stretch-receptor feedback with information about the length of the region posterior to the SMD motoneuron. Furthermore, the co-existence of both mechanisms in the worm would be feasible.

Our model integrates the head motorneuron circuit and the VNC motorneuron circuit within a physical model of the body and environment, such that the forward motion of the model emerges from the undulation of the body. Although a number of computational models had considered the head motorneuron circuit in the absence of a physical framework of the body and environmental forces [49, 54, 55, 56], previous neuro-mechanical models of forward locomotion had either assumed an oscillator in the head [14] or modeled the head circuit as an additional VNC unit [10]. The reflexive pattern generator hypothesis for oscillations in the head circuit that emerged from our evolutionary experiments had been considered in two previous models [49, 54]. We highlight here the most substantive differences between these two previous models and the one proposed here. First, in previous models the circuit responsible for oscillations included a broad range of head interneurons and motorneurons. In the Sakata and Shingai model [54], these classes include AIB, AIZ, AVB, AVK, RIA, RIB, RIC, RIG, RIS, RIV, RMD, RME, SAA, SMB, and SMD. In the Karbowski et al. model [49], the neurons were identified more abstractly as one of several possible head interneurons subsets, including AIZ, AIA, AWA, and AIZ or RIB, RIG, URY, and RIB, SAA, and head motorneurons including one of either SMB or SMD, and RME. In contrast, in the current model we demonstrate that a minimal set of head motorneurons (specifically SMD and RME) are sufficient to generate oscillations. Second, in the previous models the stretch receptor feedback into the head interneurons was postulated to come from SAA and was thus modeled to receive stretch information from the head posture. In contrast, the current model postulates that stretch receptor feedback from SMD is sufficient to drive oscillations in the head using postural information from regions in the head and posterior to the head. Third, in the previous models the oscillations in the head circuit were imposed downstream premotor command interneurons (e.g., AVB and PVC), which were then communicated to VNC motorneurons. However, the activity of these neurons has since been demonstrated not to correlate with locomotion undulations [27, 29]. In contrast, in the current model we demonstrate that the oscillations in the head motorneurons can be propagated to the VNC motorneurons through stretch-receptor feedback. Finally, in previous models the parameters of the head circuit were hand-designed to generate oscillations. In the current model, we do not assume that oscillations can only be generated in the head; oscillations in the head emerge from the evolutionary optimization process given the neuroanatomical constraints.

We have demonstrated that a neuro-mechanical model of the worm with short-range and posteriorly directed proprioceptive feedback in the VNC is sufficient to propagate the wave along the body and produce forward locomotion. A key component in our model is that we allow the stretch receptor conductance to generate a depolarizing response to compression and a polarizing response to stretch, relative to the local segment resting length, in line with recent findings [6]. A detailed analysis of the solutions revealed five key mechanisms for sustaining the proprioceptively driven dorsoventral undulations in the model. (a) The dorsoventral undulation generated in the head motoneuron circuit is propagated posteriorly to the VNC, despite the lack of direct synapses between the head motoneurons and VNC motoneurons, through stretch-receptor feedback from the anterior-most VNC neural unit. (b) The wave is propagated along the rest of the VNC neuromuscular units primarily through stretch-receptor feedback from the region immediately anterior to it. (c) Bistable motoneurons are not necessary for sustaining the proprioceptively driven dorsoventral undulations in the model. (d) Despite the role of stretch-receptor feedback, the inclusion of a biomechanical model revealed that the passive mechanics of the body play a substantial role in the propagation of the undulation, in the absence of entire subregions of the VNC. (e) The contribution from the inter-unit gap junctions was relatively minor, serving mostly to dampen curvature. The proposed model is consistent with the recent findings that in the absence of AVB-B gap junction inputs driving B-class motoneurons to intrinsic oscillatory activity, proprioceptive couplings can still propagate bending waves throughout the majority of the length of the body [57]. All of these postulated mechanisms would be promising to investigate further experimentally.

Despite the breadth of knowledge about the neurobiology, anatomy and physics of *C. elegans*, there are still a number of unanswered questions about the neuromechanical basis of one of its most basic behaviors. Our model proposes a head motorneuron circuit that can generate oscillations and a VNC motoneuron circuit that can propagate the wave using stretch-receptor feedback in a mechanical model of the body, altogether sufficient to propel the worm forward in agar. Furthermore, we demonstrate a methodology to systematically explore different mechanisms that match behavior given biological assumptions. Further work will involve matching the behavior of the integrated neuromechanical model to the effect produced from optogenetic and physical manipulations reported in recent experiments [57, 58]. Ultimately, improving our understanding of forward locomotion will allow us to study more complex behaviors that may require contributions from additional neural circuits.

## Acknowledgments

The work in this paper was supported in part by NSF grant IIS-1524647.

## References

1. JG White, E Southgate, JN Thomson, and S Brenner. The structure of the nervous system of the nematode Caenorhabditis elegans. Philos Trans R Soc Lond B Biol Sci, 275(938):327–348, 1986.

2. RH Waterston. Muscle. In WB Wood, editor, The nematode C. elegans, pages 281–335. Cold Spring Harbor Laboratory Press, New York, 1988.

3. G Haspel and MJ O’Donovan. A perimotor framework reveals functional segmentation in the motoneuronal network controlling locomotion in *Caenorhabditis elegans*. J Neurosci, 31(41): 14611–14623, 2011.

4. P Babington. C. elegans II. Cold Spring Harbour Laboratory Press, New York, 2 edition, 1997.

5. N Tavernarakis, W Shreffler, S Wang, and M Driscoll. unc-8, a deg/enac family member, encodes a subunit of a candidate mechanically gated channel that modulates *C. elegans* locomotion. Neuron, 18:107–119, 1997.

6. Q Wen, MD Po, E Hulme, S Chen, X Liu, SW Kwok, M Gershow, AM Leifer, V Butler, C Fang-Yen, et al. Proprioceptive coupling within motor neurons drives *C. elegans* forward locomotion. Neuron, 76(4):750–761, 2012.

7. J Gjorgjieva, D Biron, and G Haspel. Neurobiology of *Caenorhabditis elegans* locomotion: where do we stand? Bioscience, 64(6):476–486, 2014.

8. N Cohen and T Sanders. Nematode locomotion: dissecting the neuronal–environmental loop. Curr Opin Neurobiol, 25:99–106, 2014.

9. M Zhen and ADT Samuel. *C. elegans* locomotion: small circuits, complex functions. Curr Opin Neurobiol, 33:117–126, 2015.

10. JH Boyle, S Berri, and N Cohen. Gait modulation in *C. elegans*: an integrated neuromechanical model. Front Comput Neurosci, 6:10, 2012. doi:10.3389/fncom.2012.00010.

11. S Berri, JH Boyle, M Tassieri, IA Hope, and N Cohen. Forward locomotion of the nematode *C. elegans* is achieved through modulation of a single gait. HFSP J, 3(3):186–193, 2009.

12. JH Boyle. C. elegans locomotion: An integrated approach. PhD thesis, University of Leeds, 2010.

13. N Cohen and JH Boyle. Swimming at low reynolds number: a beginner’s guide to undulatory locomotion. Contemporary Physics, 51:103–123, 2010.

14. E Niebur and P Erdös. Theory of the locomotion of nematodes: Dynamics of undulatory progression on a surface. Biophys J, 60(5):1132–1146, 1991.

15. J Karbowski, CJ Cronin, A Seah, JE Mendel, D Cleary, and PW Sternberg. Conservation rules, their breakdown, and optimality in *Caenorhabditis* sinusoidal locomotion. Journal of Theoretical Biology, 242(3):652–669, 2006. ISSN 0022-5193. doi:10.1016/j.jtbi.2006.04.012.

16. P Sauvage, M Argentina, J Drappier, T Senden, J. SimÃl’on, and JM Di Meglio. An elasto-hydrodynamical model of friction for the locomotion of *Caenorhabditis elegans*. Journal of Biomechanics, 44(6):1117–1122, 2011. ISSN 0021-9290. doi:10.1016/j.jbiomech.2011.01.026.

17. R Rabets, M Backholm, K Dalnoki-Veress, and Ryu WS. Direct measurements of drag forces in *C. elegans* crawling locomotion. Biophysical Journal, 107(8):1980–1987, 2014. ISSN 0006-3495. doi:10.1016/j.bpj.2014.09.006.

18. J Lighthlil. Flagellar hydrodynamics. SIAM Rev, 18:161–230, 1976.

19. HR Wallace. Wave formation by infective larvae of the plant parasitic nematode *Meloidogyne javanica*. Nematologica, 15(1):65–75, 1969.

20. P Sauvage. Etude de la locomotion chez C. elegans et perturbations mecaniques du mouvement. PhD thesis, Universite Paris, 2007.

21. ZF Altun and DH Hall. Muscle system, somatic muscle. In WormAtlas. 2009. doi:10.3908/wormatlas.1.7.

22. AV Hill. The heat of shortening and the dynamics constants of muscle. Proc. R. Soc. London B, 126:136–195, 1938.

23. B Milligan, N Curtin, and Q Bone. Contractile properties of obliquely striated muscle from the mantle of squid (*Alloteuthis subulata*) and cuttlefish (*Sepia officinalis*). J Exp Biol, 200(Pt 18): 2425–24236, 1997.

24. C, AM Leifer Fang-Yen, M Gershow, MJ Alkema, and ADT Samuel. Optogenetic manipulation of neural activity in freely moving *Caenorhabditis elegans*. Nature Methods, 8 147–152, 2011. doi:10.1038/nmeth.1554.

25. JB Rand and ML Nonet. Neurotransmitter assignments for specific neurons. In C. elegans II, pages 1049–1052. Cold Spring Harbour Laboratory Press, New York, 1997.

26. M Chalfie, JE Sulston, JG White, E Southgate, JN Thomson, and S Brenner. The neural circuit for touch sensitivity in *Caenorhabditis elegans*. J Neurosci, 5(4):956–964, 1985.

27. S Faumont, G Rondeau, TR Thiele, KJ Lawton, KE McCormick, M Sottile, O Griesbeck, ES Heckscher, WM Roberts, Doe CQ, et al. An image-free opto-mechanical system for creating virtual environments and imaging neuronal activity in freely moving *Caenorhabditis elegans*. PLoS One, 6(9):e24666, 2011. doi:10.1371/journal.pone.0024666.

28. G Haspel, MJ O’Donovan, and AC Hart. Motoneurons dedicated to either forward or backward locomotion in the nematode *Caenorhabditis elegans*. J Neurosci, 30(33):11151–11156, 2010.

29. T Kawano, Po MD, S Gao, G Leung, WS Ryu, and M Zhen. An imbalancing act: gap junctions reduce the backward motor circuit activity to bias *C. elegans* for forward locomotion. Neuron, 72(4):572–586, 2011.

30. LR Varshney, BL Chen, E Paniagua, DH Hall, and DB Chklovskii. Structural properties of the *Caenorhabditis elegans* neuronal network. PLoS Comput Biol, 7(2):e1001066, 2011. doi:10.1371/ journal.pcbi.1001066.

31. SL McIntire, E Jorgensen, J Kaplan, and HR Horvitz. The gabaergic nervous system of *Caenorhabditis elegans*. Nature, 364(6435):337–41, 1993.

32. SR Lockery and MB Goodman. The quest for action potentials in *C. elegans* neurons hits a plateau. Nat Neurosci, 12(4):377–378, 2009.

33. JE Mellem, PJ Brockie, DM Madsen, and AV Maricq. Action potentials contribute to neuronal signaling in *C. elegans*. Nat Neurosci, 11(8):865–867, 2008.

34. EJ Izquierdo and SR Lockery. Evolution and analysis of minimal neural circuits for klinotaxis in *Caenorhabditis elegans*. J Neurosci, 30(39):12908–12917, 2010.

35. EJ Izquierdo and RD Beer. Connecting a connectome to behavior: an ensemble of neuroanatomical models of *C. elegans* klinotaxis. PLoS Comput Biol, 9(2):e1002890, 2013. doi:10.1371/journal.pcbi.1002890.

36. M Kuramochi and M Doi. A computational model based on multi-regional calcium imaging represents the spatio-temporal dynamics in a *Caenorhabditis elegans* sensory neuron. PLoS One, 12(1):e0168415, 2017. doi:10.1371/journal.pone.0168415.

37. TH Lindsay, TR Thiele, and SR Lockery. Optogenetic analysis of synaptic transmission in the central nervous system of the nematode *Caenorhabditis elegans*. Nat Commun, 2(306), 2011. doi:10.1038/ncomms1304.

38. SR Wicks, CJ Roehrig, and CH Rankin. A dynamic network simulation of the nematode tap withdrawal circuit: predictions concerning synaptic function using behavioral criteria. J Neurosci, 16(12):4017–4031, 1996.

39. JM Kunert, JL Proctor, SL Brunton, and JN Kutz. Spatiotemporal feedback and network structure drive and encode *Caenorhabditis elegans* locomotion. PLoS Comput Biol, 13(1):e1005303, 2017. doi:10.1371/journal.pcbi.1005303.

40. EO Olivares, EJ Izquierdo, and RD Beer. Potential role of a ventral nerve cord central pattern generator in forward and backward locomotion in *Caenorhabditis elegans*. Network Neuroscience, 0(a):1–32, 2017. doi:10.1162/NETN\_a\_00036.

41. RD Beer. On the dynamics of small continuous-time recurrent neural networks. Adapt Behav, 3(4):469–509, 1995.

42. W Li, Z Feng, PW Sternberg, and Xu XZS. A *C. elegans* stretch receptor neuron revealed by a mechanosensitive trp channel homologue. Nature, 440:684–687, 2006.

43. WR Schafer. Proprioception: a channel for body sense in the worm. Curr. Biol., 16:R509–R511, 2006.

44. XN Shen, J Sznitman, P Krajacic, T Lamitina, and PE Arratia. Undulatory locomotion of *Caenorhabditis elegans* on wet surfaces. Biophysical Journal, 102(12):2772–2781, 2012. doi:10.1016/j.bpj.2012.05.012.

45. DT Omura, DA Clark, ADT Samuel, and HR Horvitz. Dopamine signaling is essential for precise rates of locomotion by *C. elegans*. PLOS ONE, 7(6):1–9, 06 2012. doi:10.1371/journal. pone.0038649.

46. E Cohen, E Yemini, W Schafer, DG Feitelson, and M Treinin. Locomotion analysis identifies roles of mechanosensory neurons in governing locomotion dynamics of *C. elegans*. Journal of Experimental Biology, 215(20):3639–3648, 2012. doi:10.1242/jeb.075416.

47. CJ Cronin, JE Mendel, S Mukhtar, Kim YM, RC Stirbl, J Bruck, and PW Sternberg. An automated system for measuring parameters of nematode sinusoidal movement. BMC Genetics, 6(5), 2005. doi:10.1186/1471-2156-6-5.

48. C Fang-Yen, M Wyart, J Xie, R Kawai, T Kodger, S Chen, and ADT Samuel. Biomechanical analysis of gait adaptation in the nematode *Caenorhabditis elegans*. Proceedings of the National Academy of Sciences of the United States of America, 107(47):20323–20328, 2010. doi:10.1073/pnas. 1003016107.

49. J Karbowski, G Schindelman, CJ Cronin, A Seah, and PW Sternberg. Systems level circuit model of *C. elegans* undulatory locomotion: mathematical modeling and molecular genetics. J Comput Neurosci, 24(3):253–276, 2008.

50. JT Pierce-Shimomura, BL Chen, JJ Mun, R Ho, R Sarkis, and SL McIntire. Genetic analysis of crawling and swimming locomotory patterns in *C. elegans*. Proceedings of the National Academy of Sciences of the United States of America, 105(52):20982–20987, 2008. doi:10.1073/ pnas.0810359105.

51. E Yemini, T Jucikas, LJ Grundy, AEX Brown, and WR Schafer. A database of *C. elegans* behavioral phenotypes. Nature Methods, 10(9):877–879, 2013. doi:10.1038/nmeth.2560.

52. A Vidal-Gadea, S Topper, L Young, A Crisp, L Kressin, E Elbel, and JT Pierce-Shimomura. *Caenorhabditis elegans* selects distinct crawling and swimming gaits via dopamine and serotonin. Proceedings of the National Academy of Sciences of the United States of America, 108 (42):17504–17509, 2011. doi:10.1038/nmeth.2560.

53. VJ Butler, R Branicky, E Yemini, JF Liewald, A Gottschalk, RA Kerr, DB Chklovskii, and WR Schafer. A consistent muscle activation strategy underlies crawling and swimming in *Caenorhabditis elegans*. Journal of The Royal Society Interface, 12(102), 2015. doi:10.1098/rsif.2014. 0963.

54. K Sakata and R Shingai. Neural network model to generate head swing in locomotion of *Caenorhabditis elegans*. Network, 15(3):199–216, 2004.

55. X Deng and Xu JX. A 3d undulatory locomotion model inspired by *C. Elegans* through DNN approach. Neurocomput., 131:248–264, May 2014. ISSN 0925-2312. doi:10.1016/j.neucom.2013. 10.019.

56. X Deng, Xu JX, J Wang, G Wang, and Q Chen. Biological modeling the undulatory locomotion of *C. elegans* using dynamic neural network approach. Neurocomputing, 186:207–217, 2016.

57. T Xu, J Huo, S Shao, M Po, T Kawano, Y Lu, M Wu, M Zhen, and Q Wen. Descending pathway facilitates undulatory wave propagation in *Caenorhabditis elegans* through gap junctions. Proceedings of the National Academy of Sciences, 2018. ISSN 0027-8424. doi:10.1073/pnas. 1717022115.

58. AD Fouad, S Teng, JR Mark, S Liu, P Alvarez-Illera, Ji H, A Du, PD Bhirgoo, E Cornblath, SA Guan, and C Fang-Yen. Distributed rhythm generators underlie *Caenorhabditis elegans* forward locomotion. eLife, 7:e29913, 2018. doi:10.7554/eLife.29913.

